# Ancestry-Linked IL-10 Signaling and Macrophage Activation Modulate Fibroblast Responses to Oxidative Stress in a PEG-Based Microphysiological System

**DOI:** 10.64898/2026.05.04.722732

**Authors:** Nana Owusu-Boaitey, Alison M. Veintimilla, Miriam Tamaño-Blanco, Paul Parodi, Katriel Barcellano Kasayan, Sanjana Ranasinghe, Erika Moore

## Abstract

Ancestry-associated immune differences influence fibrosis risk, however, how fibrosis-associated pathways vary across individuals remains poorly understood. Fibroblasts are a main cell type involved in fibrosis. The fibroblast response is shaped by cytokine signaling and macrophage activation. The extent to which these pathways vary across individuals, and how ancestry-associated immune differences influence fibrosis risk, remains poorly understood. Here, a poly(ethylene glycol) (PEG)-based hydrogel microphysiological system was leveraged to model fibroblast–macrophage interactions following oxidative stress and to integrate donor-specific immune signals using matched macrophages and serum. Individuals of self-reported African ancestry exhibited higher monocyte expression of CCL4, lower monocyte expression of OXER1, and increased serum IL-10, compared to individuals of European ancestry. Within the hydrogel, oxidative stress reduced fibroblast prevalence while inducing Ki67 and p16. Exogenous TGF-β1 increased fibroblast prevalence and collagen 3 production but did not independently increase α-SMA. Incorporating donor-specific macrophages and serum revealed that cultures from individuals of European ancestry demonstrated higher fibroblast α-SMA and p16 expression. Pharmacologic inhibition of IL-10 further increased α-SMA, particularly in African ancestry–derived cultures, identifying IL-10 as a key protective signal limiting fibroblast activation. This hydrogel system provides a platform for dissecting inter-individual immune variation and identifying mechanisms underlying ancestry-associated fibrosis risk.

## Introduction

Soft-tissue wound repair depends on the coordinated recovery of damaged cells and the reconstitution of the extracellular matrix (ECM).^1^ Following injury, fibroblasts proliferate to replace damaged cells and deposit ECM proteins that restore tissue structure.^2,3^ When ECM deposition is dysregulated, the repair response progresses toward fibrosis. During fibrosis, fibroblasts proliferate and deposit excessive levels of ECM proteins.^4^ This excess ECM deposition results in the failure to recapitulate normal tissue function.^4^ Over 100 million people each year experience dermal fibrosis in the form of excessive scarring.^5^ Understanding the cellular and molecular cues that regulate fibroblast behavior after injury is critical for identifying strategies to prevent or treat fibrosis.

*In vitro* fibrosis models improve understanding of the mechanisms driving fibroblast responses to injury. These models seek to replicate responses observed *in vivo*, including fibroblast expression of the activation marker alpha-smooth muscle actin (α-SMA) and fibroblast deposition of ECM proteins including collagen 3.^6,7^ Fibroblast proliferation during repair is also associated with expression of the nuclear protein Ki67 in these model systems and *in vivo*.^8,9^

Oxidative provides an effective means of inducing and modeling fibroblast injury. Oxidative stress induced sterile, chemical injury in *in vitro* models.^2,3,10,11^ Following oxidative stress, fibroblasts increase expression of fibrosis-associated genes.^3^ The cyclin-dependent kinase inhibitor p16 correlates with oxidative stress and senescence in profibrotic contexts.^8,9,12^ Fibroblast response can be modulated using serum from patients with fibrotic conditions to assess the role of soluble factors.^13^

Cytokine signaling is a key regulator of fibroblast responses and fibrosis risk. Macrophage immune cells and their precursor, monocytes, are important sources of fibroblast modulating cytokines, such as TGF-β and IL-10. The pleiotropic cytokine transforming growth factor-β1 (TGF-β1) is well-established as a driver of fibroblast survival and profibrotic ECM production, including collagen deposition.^4,14^ Oxidative stress induces TGF-β signaling pathways.^3,15^ Inhibition of macrophage recruitment or activation reduces TGF-β levels and mitigates fibrosis in several disease models.^16–19^ In parallel, interleukin-10 (IL-10) can constrain excessive inflammation and has been implicated in limiting fibrosis.^20,21^ IL-10 further skews macrophages towards an anti-inflammatory phenotype.^22^ How macrophage behavior and cytokine profiles jointly shape fibroblast survival, activation, and ECM deposition following injury-associated oxidative stress remains incompletely understood.

Demographic factors, including ancestry, contribute to fibrosis risk. Self-identified Black individuals have demonstrated greater serum levels of inflammatory markers.^23,24^ African ancestry has been correlated with stronger macrophage inflammatory response.^25^ This suggests that ancestry-associated differences in inflammatory pathways may influence susceptibility to fibrosis.^26,27^ However, there are limited in vitro models into how ancestry-associated differences in macrophage function and circulating cytokines affect fibroblast responses to injury.

Experimental models that integrate ancestry-linked immune and serum factors with fibroblast behavior could therefore help elucidate pathways that modulate fibrosis risk.

Biomaterial-based models offer a controlled platform to interrogate fibrotic responses.^7,28^ Hydrogels can mimic key aspects of the three-dimensional ECM while allowing independent modulation of cell composition, cytokine signaling, and injury cues.^29^ Poly(ethylene glycol) (PEG)-based hydrogels provide a tunable environment to study fibroblast–macrophage interactions and quantify injury-induced changes in proliferation, activation, and ECM composition.^30,31^

Utilizing a functionalized PEG-based system we have developed an *in vitro* 3D model capable of sustaining fibroblasts following oxidative stress. We used this model to study ancestry-associated differences in fibroblast response and activation amongst individuals of African or European ancestry. We further assessed the role of anti-inflammatory macrophage activation, IL-10, and TGF-β1 signaling in the fibroblast response to oxidative stress. We hypothesized that IL-10 signaling inhibits fibroblast activation in an ancestry-dependent manner following injury. Anti-inflammatory activation and TGF-β1 signaling would promote fibroblast activation and ECM deposition. Together, these results validate a 3D hydrogel model elucidating mechanisms underlying the fibroblast response to sterile injury.

## Results

### Transcriptomic analysis of monocytes and proteomic analysis of serum from participants of different ancestry

To assess correlations between self-reported ancestry and inflammatory signaling, monocyte gene expression and serum cytokines were analyzed for individuals of self-reported African or European ancestry (Fig. 1a). Ancestry classification was based on participant self-report on a demographic survey.^23,24^ Gene expression analysis revealed a 4.3-fold increase (p < 0.05) in expression of the chemokine CCL4 for individuals of African ancestry compared to those of European ancestry (Fig. 1b and 1c). Individuals of European ancestry had a 1.6-fold increase (p < 0.05) in expression of OXER1, a receptor associated with monocyte migration (Fig. 1b and 1c). Mass spectrometry of serum proteins isolated from the same participants showed a 2.3-fold increase (p < 0.05) in circulating IL-10 for individuals of European ancestry (Fig. 1d). Together, these ancestry-associated differences in our cohort of individuals highlight differences in monocyte inflammatory signaling and serum IL-10 which motivates the need to consider ancestry in our hydrogel model.

**Figure 1.**
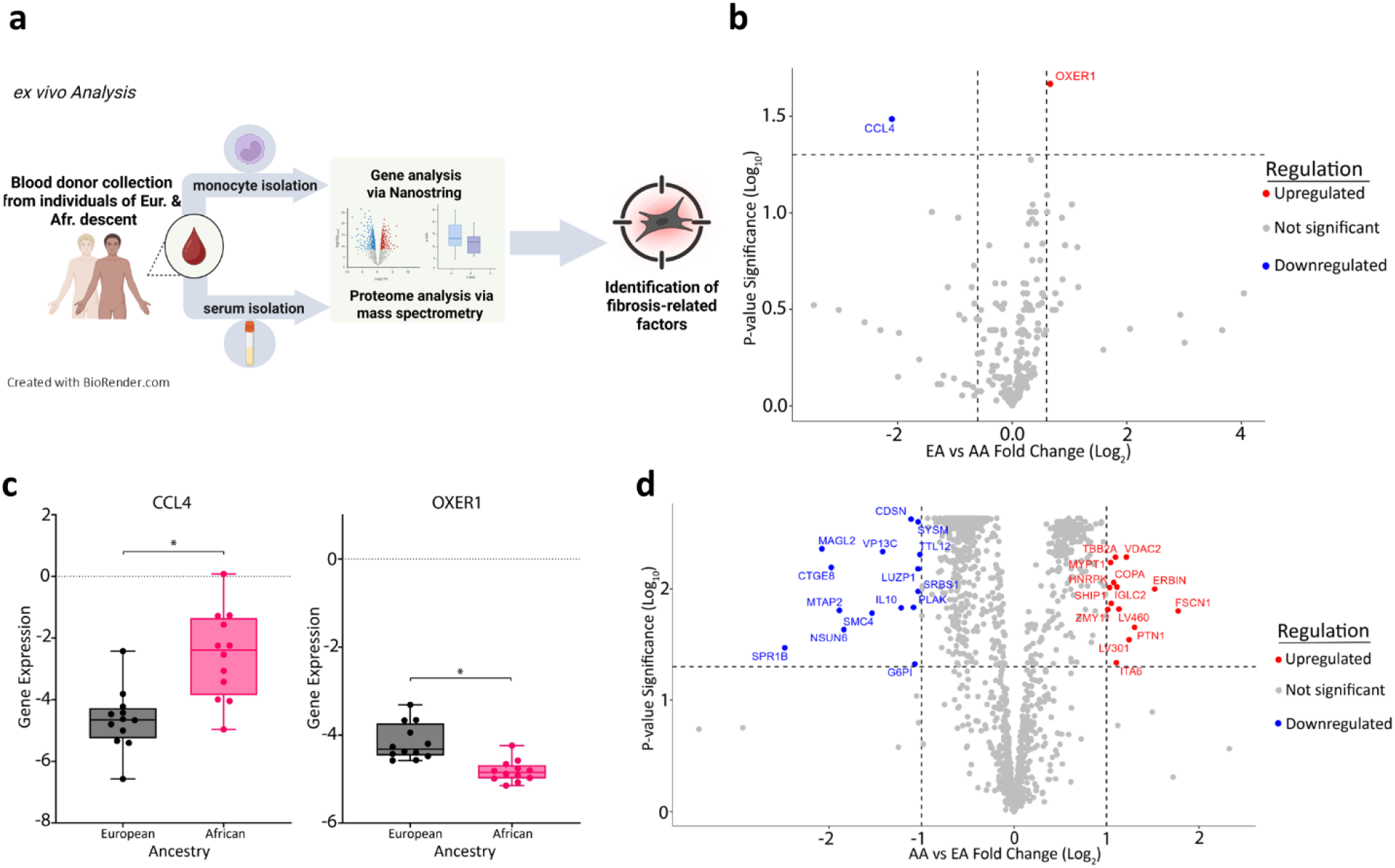
(a) *Ex vivo* assessment of macrophage and soluble factors in injury. (b), (c) Gene expression for recruited participants of European ancestry (EA) and African ancestry (AA) based on NanoString. Fold change represents European vs. African expression and is log transformed. n = 16 biological replicates per ancestry group. (d) Serum protein levels and for recruited participants of European ancestry and African ancestry based on mass spectroscopy. Fold change represents African vs. European protein levels and is log transformed. n = 16 biological replicates per ancestry group. \**p*-value < 0.05. Created in BioRender. Moore, E. (2026) https://BioRender.com/6neh9dq.

### Anti-inflammatory macrophage activation and TGF-β1 signaling promote fibrosis-associated fibroblast responses in a hydrogel model of sterile injury

To establish the oxidative stress conditions within our hydrogel model used in subsequent experiments, dermal fibroblasts encapsulated in PEG hydrogels were treated for 24 hours with 0, 125, or 250 μM of H_2_O_2_, followed by recovery for 0, 3, or 7 days (Fig. 2a). The hydrogel provided sites for cell adhesion through inclusion of the peptide arginine-glycine-aspartate-serine (RGDS). Encapsulated cells were able to remodel the hydrogel matrix environment due to the presence of PQ (GGGPQGIWGQGK) peptide.^30,31^ We selected PEG concentrations that resulted in hydrogel stiffness similar to that of the skin.^32–34^ After 7 days, fibroblast prevalence decreased by approximately 2-fold following 125 μM H_2_O_2_ (p < 0.01), and by approximately 6-fold following 250 μM (p < 0.001) (Fig. 2b; Supp. Fig. 1a). At 7 days the proportion of fibroblasts positive for the protein Ki67 increased from 21% to 44% for 0 μM H_2_O_2_ treatment compared to 125 μM (p < 0.01). Fibroblast expression of p16 increased from 28% to 64% (p < 0.001) (Fig. 2b and 2c; Supp. Fig. 1b and 1c). Based on these results, 125 μM H_2_O_2_ was selected for subsequent experiments, as it reduced fibroblast prevalence while maintaining measurable proliferative and stress-associated responses.

**Figure 2.**
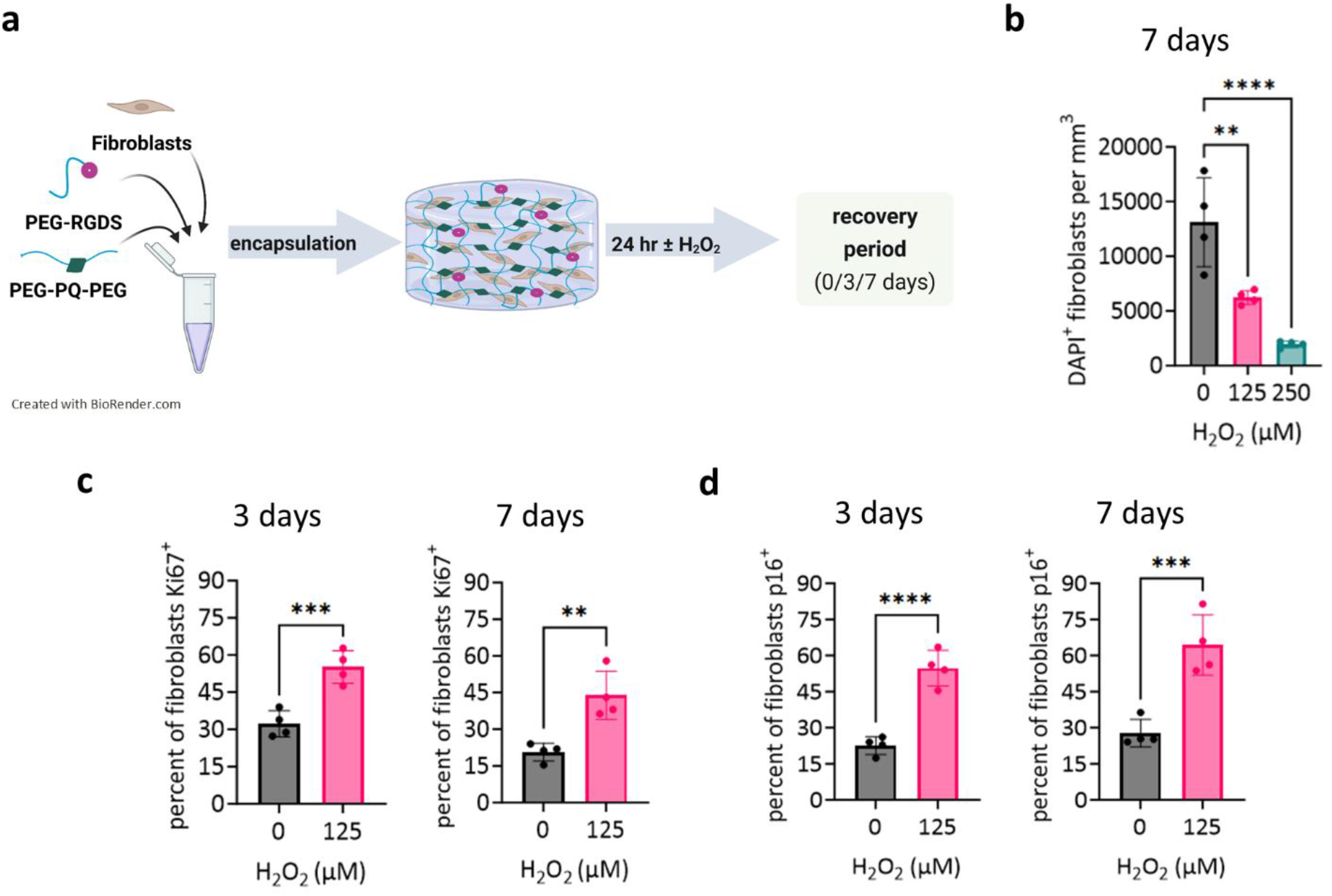
Oxidative stress reduces fibroblast survival and induces proliferative and stress-associated markers in encapsulated 3D hydrogel. (a) Encapsulation of fibroblasts in PEG-based hydrogel, followed by oxidative stress and recovery. (b) Number of fibroblasts per volume of hydrogel at 0, 3, and 7 days after H_2_O_2_ treatment. (c) Percentage of fibroblasts expressing Ki67, and (d) p16 at 3 and 7 days after H_2_O_2_ treatment. n = 4 biological replicates. Data shown as mean ± standard deviation. The Kruskal-Wallis test was used to calculate p-values and statistical significance. \*\**p*-value < 0.01; \*\*\**p*-value < 0.001, \*\*\*\**p*-value < 0.0001. Created in BioRender. Moore, E. (2026) https://BioRender.com/6neh9dq.

To determine the role of macrophages in fibroblast activation, fibroblasts were co-encapsulated with unactivated, inflammatory, or anti-inflammatory macrophages. This was followed by culture for 7 days without H_2_O_2_ (Fig. 3a). By 7 days after oxidative stress, anti-inflammatory macrophages increased fibroblast α-SMA expression from 14% to 24% (p < 0.001) (Fig. 3b). Fibroblast expression of p16 and intracellular collagen 3 did not significantly change (Fig. 3c and 3d).

**Figure 3.**
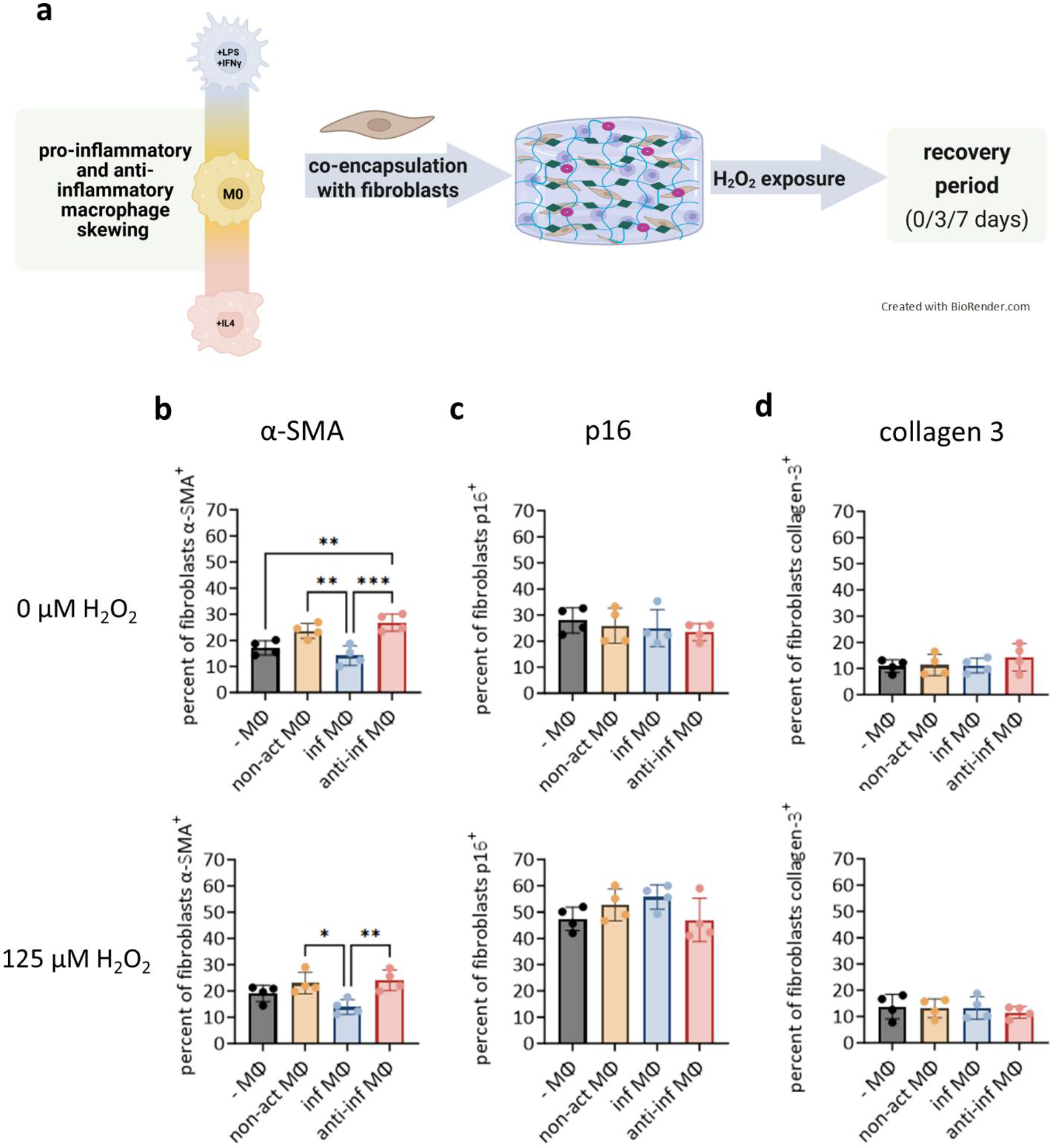
Anti-inflammatory macrophages induce fibroblast activation markers in the absence of donor-specific immune signals. (b) Number of fibroblasts expressing alpha smooth muscle actin [α-SMA], (c) p16, and (d) intracellular collagen 3 by immunocytochemistry at 7 days after H_2_O_2_ treatment and encapsulated without macrophages (-Mθ), unactivated macrophages (non-act Mθ), inflammatory macrophages (inf Mθ), and anti-inflammatory macrophages (anti-inf Mθ). n = 4 biological replicates for macrophage donors. Data shown as mean ± standard deviation. The Kruskal-Wallis test was used to calculate p-values and statistical significance. Data shown as mean ± standard deviation. \**p*-value < 0.05; \*\**p*-value < 0.01; \*\*\**p*-value < 0.001. Created in BioRender. Moore, E. (2026) https://BioRender.com/6neh9dq.

We next assessed the effects of TGF-β signaling on fibroblast responses to oxidative stress. Fibroblasts were cultured for 24 hours with or without TGF-β1, with or without H_2_O_2_. This was followed by culture for 7 days without H_2_O_2_. At 7 days following oxidative stress, TGF-β1 increased fibroblast prevalence by 1.5-fold (p < 0.01) (Fig. 4a) and increased Ki67+ fibroblasts by 12% (p < 0.01) (Fig. 4b). TGF-β1 treatment did not significantly change p16 or α-SMA (Fig. 4c and 4d). TGF-β1 treatment increased collagen 3 expression from 15% to 29% (p < 0.0001) (Fig. 4e). These results suggest a role for TGF-β1 in supporting fibroblast survival and ECM production after oxidative stress.

**Figure 4.**
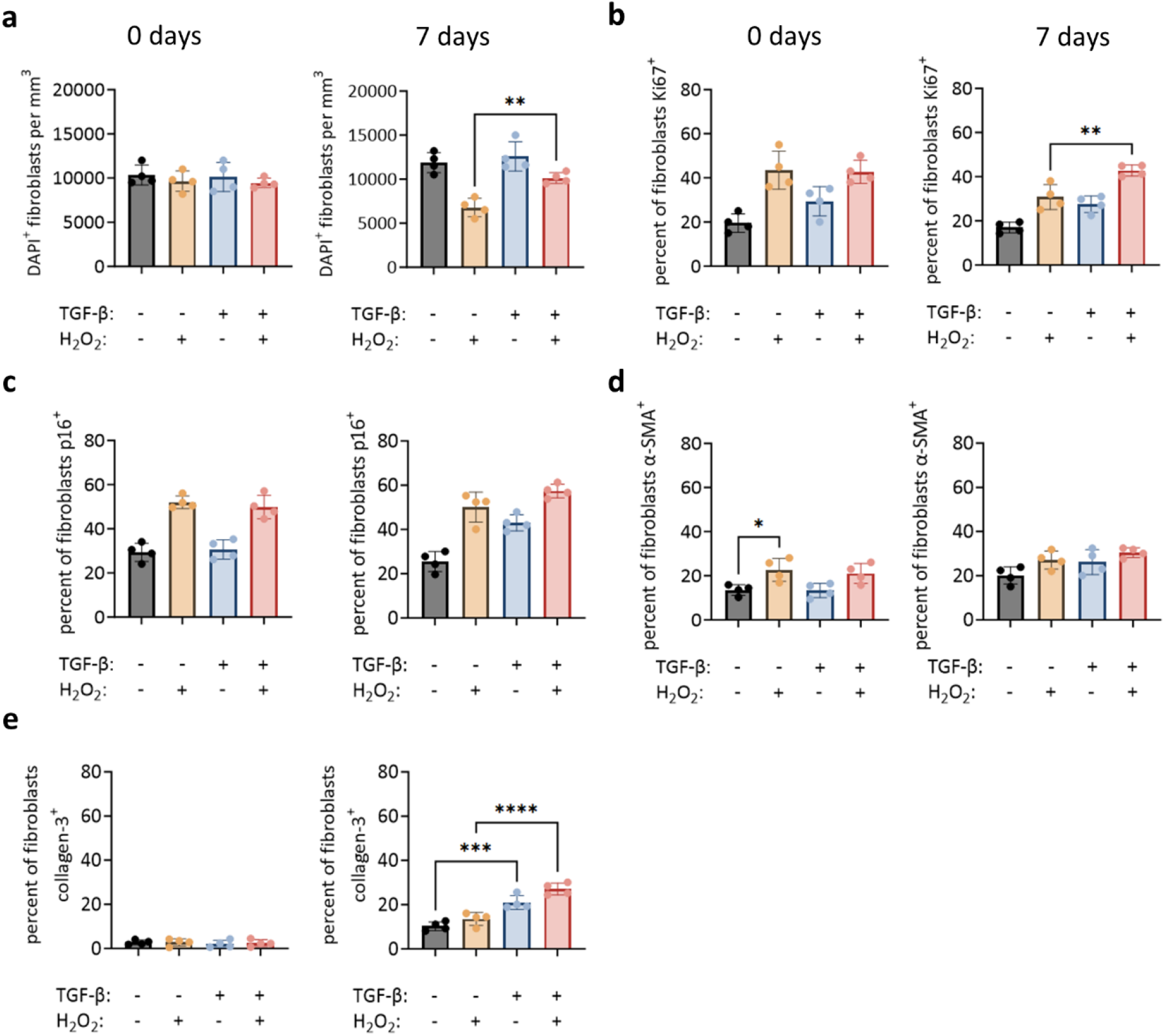
TGF-β signaling enhances fibroblast survival and collagen production after oxidative stress but is not required for macrophage-induced fibroblast activation. (a) Number of fibroblasts per volume of hydrogel, (b) percentage of fibroblast expressing Ki67, (c) p16, (d) intracellular collagen 3, and (e) α-SMA by immunocytochemistry at 0 and 7 days after H_2_O_2_ and TGF-β1 treatment. n = 4 biological replicates. Data shown as mean ± standard deviation. The Kruskal-Wallis test was used to calculate p-values and statistical significance. \*\**p*-value < 0.01; \*\*\**p*-value < 0.001, \*\*\*\**p*-value < 0.0001.

### IL-10-dependent inhibition of fibroblast activation by macrophages and serum from donors of African ancestry

To evaluate how donor-specific immune signaling influences the repair response, serum and isolated macrophages from each group were incorporated into the hydrogel model. Donor macrophages were left unstimulated or stimulated towards inflammatory or anti-inflammatory. Each macrophage group was then co-encapsulated with fibroblasts in media containing matched donor serum. After 24 hours with or without H_2_O_2_, the cultures were allowed to recover for 7 days prior to analysis. Ancestry-associated differences were noted in the unactivated and anti-inflammatory macrophage phenotype, but not under inflammatory macrophage conditions. Fibroblasts cultured with unactivated macrophages from individuals of European ancestry exhibited higher α-SMA expression (31%) than those from African ancestry (16%, p < 0.0001) (Fig. 5a). This ancestry-associated difference persisted even in the absence of oxidative stress (25% vs. 18%, p < 0.001). Ancestry-associated differences were also observed in p16 expression. Under unactivated macrophage conditions and without oxidative stress, 11% of fibroblasts from African donors expressed p16 compared to 22% from European ancestry donors (p < 0.0001).

**Figure 5.**
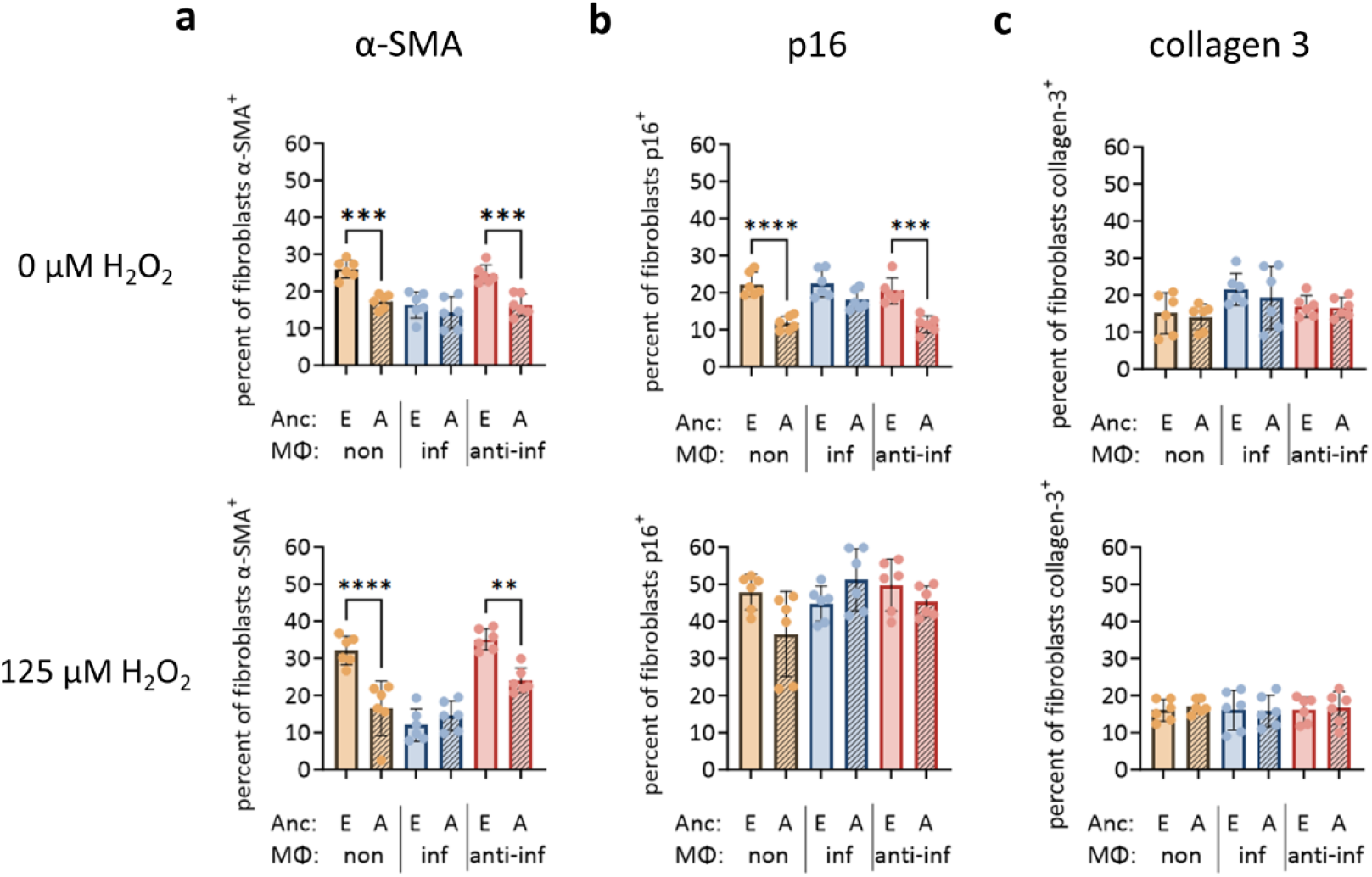
Self-reported European ancestry is associated with increased fibroblast activation and p16 expression in a macrophage–fibroblast hydrogel model. (a) Percentage of fibroblasts expressing α-SMA, (b) p16, and (c) intracellular collagen 3 by immunocytochemistry at 7 days after H_2_O_2_ treatment, culture in donor serum, and co-encapsulation with unactivated macrophages (non), inflammatory macrophages (inf), and anti-inflammatory macrophages (anti-inf). Macrophages and serum are from donors of self-reported European or African ancestry. n = 6 biological replicates per ancestry group. The Kruskal-Wallis test was used to calculate p-values and statistical significance. \*\**p*-value < 0.01; \*\*\**p*-value < 0.001, \*\*\*\**p*-value < 0.0001.

Under anti-inflammatory macrophage conditions, these fibroblast percentages were 12% vs. 19% (p < 0.001) (Fig. 5b). No significant differences were observed by ancestry for protein expression collagen 3 (Fig. 5c).

Finally, to determine whether IL-10 signaling contributed to ancestry-associated differences in fibroblast activation, IL-10 signaling was inhibited using Stattic, a STAT-3 inhibitor (Fig. 6a). Stattic treatment did not significantly affect collagen 3 or p16 expression (Supp. Fig. 2a and 2b). However, following oxidative stress, IL-10 inhibition increased α-SMA expression by approximately 1.5-fold for fibroblasts cultured with unactivated macrophages and serum from African ancestry donors, with a similar increase under anti-inflammatory macrophage conditions (Fig. 6b). No significant differences were observed with inflammatory macrophages (Fig. 6b). These findings indicate that IL-10 signaling limited fibroblast activation for African donors in our model and suggest that lower IL-10 signaling in African ancestry cultures contributed to increased α-SMA expression following oxidative stress.

**Figure 6.**
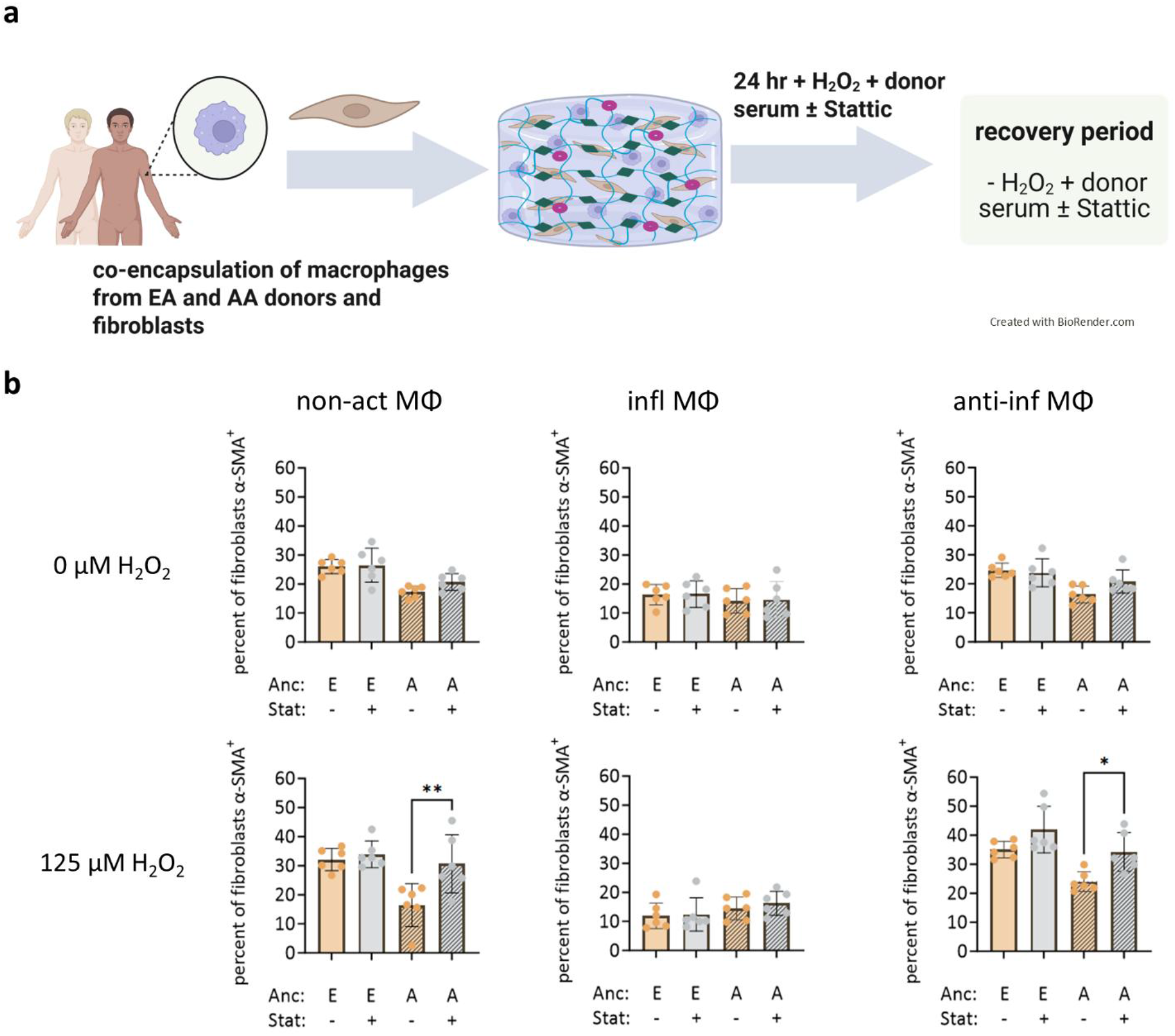
IL-10 signaling suppresses fibroblast activation, particularly in cultures from donors of African ancestry. (a) Encapsulation of fibroblasts in PEG-based hydrogel, followed by oxidative stress with culture in donor serum, with or without Stattic treatment, and co-encapsulation with macrophages, followed by a 7 day recovery period. (b) Percentage of fibroblasts expressing α-SMA after the recovery period following co-encapsulation with unactivated macrophages (non-act Mθ), inflammatory macrophages (inf Mθ), and anti-inflammatory macrophages (anti-inf Mθ). Macrophages and serum are from donors of self-reported European (E) or African (A) ancestry. n = 6 biological replicates per ancestry group. The Kruskal-Wallis test was used to calculate p-values and statistical significance. \**p*-value < 0.05; \*\**p*-value < 0.01. Created in BioRender. Moore, E. (2026) https://BioRender.com/6neh9dq.

## Discussion

In this study, we used a PEG-based hydrogel model to examine how macrophage activation, cytokine signaling, and self-reported ancestry influence fibroblast responses to oxidative stress. We integrated this *in vitro* injury model with gene expression and proteomic measurements from circulating monocytes and serum. Our results indicate that anti-inflammatory macrophage activation and TGF-β signaling support fibroblast survival and ECM production after injury. Donors of self-reported African ancestry expressed lower circulating IL-10 levels compared to European donors. In our injury model, anti-inflammatory macrophages from those of African ancestry were associated with less fibroblast activation compared to macrophages from those of European ancestry. IL-10 signaling constrained fibroblast activation for African donors but not European donors. These results further elucidate the role of ancestry, anti-inflammatory macrophage signaling, and IL-10 signaling in the fibroblast response to injury.

Our hydrogel model recapitulates several markers observed during *in vivo* tissue repair and fibrosis, while modeling skin stiffness and permitting three-dimensional cellular interactions. Following H_2_O_2_ exposure, fibroblasts exhibited increased Ki67 expression, a nuclear protein associated with fibroblast proliferation during repair.^35^ We also observed robust induction of p16, a cyclin-dependent kinase inhibitor associated with oxidative stress and senescence in profibrotic contexts.^3,9^ Approximately 60% of fibroblasts expressed p16 at later time points in our model, compared to ∼35% of dermal cells reported in mature keloid scars^9^ and less than 10% during earlier phases of the repair response.^8,12^ This suggests that our system may capture an earlier or more acute phase of the response to injury. Oxidative stress from hydrogen peroxide treatment may have also increased p16 expression. In parallel, we detected increased α-SMA expression, indicating fibroblast activation toward a myofibroblast-like phenotype.^9^ Approximately 40% of fibroblasts expressed α-SMA following oxidative stress, similar to the α-SMA expression observed in fibrotic scars.^9^ Together, these features support the use of this hydrogel platform to study fibrosis-associated fibroblast states.

Exogenous TGF-β1 increased the prevalence of fibroblasts and collagen 3 expression following oxidative stress, consistent with prior work.^3,4,15^ Interestingly, TGF-β1 did not significantly increase α-SMA in our system, in contrast to models where TGF-β robustly drives myofibroblast differentiation.^36^ This suggests that TGF-β1 alone is sufficient to support survival and ECM deposition, but additional cues may be required to fully induce α-SMA in this 3D context. One such cue is likely IL-4-induced anti-inflammatory macrophages which increased fibroblast α-SMA in our model system.

Our data highlight IL-10 as a pathway for macrophage activation to modulate fibroblast activation, consistent with prior literature on the role of IL-10 in the repair response.^37,38^ Donors of self-reported African ancestry had higher serum IL-10 levels compared to donors of European ancestry, along with increased expression of *CCL4* and *OXER1*, markers associated with increased migration to sites of inflammation.^39,40^ In the hydrogel model, fibroblasts cultured with macrophages and serum from European donors exhibited higher α-SMA and p16 expression than those cultured with serum and macrophages from African donors, particularly in the presence of oxidative stress. Pharmacological inhibition of IL-10 signaling with Stattic further increased α-SMA after injury. This increase was especially large in cultures with unactivated or anti-inflammatory macrophages from African donors. These findings support a model in which IL-10 acts as a protective anti-fibrotic signal that limits fibroblast activation. Lower IL-10 in some donor contexts may permit greater activation after injury.

Several limitations should be considered in this work. First, our macrophages were derived from circulating monocytes and polarized *in vitro*, and thus may not fully capture the diversity of macrophage subsets present in specific tissues.^41^ With respect to macrophage activation, IL-4– driven anti-inflammatory macrophages are only one of several phenotypes that can arise during repair. Second, we modeled injury using oxidative stress in a sterile context; responses to mechanical wounding or infection may involve additional pathogen-driven inflammatory signals that alter the balance between pro- and anti-fibrotic pathways.^42^ Cellular stress from hydrogel encapsulation may have also increased p16 expression, in addition to oxidative stress from hydrogen peroxide exposure. Third, we focused on a 7-day recovery window, which may not be sufficient to fully capture long-term ECM remodeling and chronic fibrosis. Extending the time course and incorporating additional injury modalities will be important future directions. Fourth, our analysis included only donors aged 18 to 45 with limited serum from each donor and thus may not extend to healing responses in pediatric or elderly populations. Fifth, Stattic may have inhibited pathways other than IL-10 signaling. We used Stattic due to improved diffusion through the hydrogel when compared to neutralizing antibodies against IL-10 receptor.^43^ Future work may assess the feasibility of other neutralizing antibodies in our system. Sixth, our analysis included only adults between the ages of 18 and 45, before the onset of postmenopausal changes in hormone levels. It remains unclear if these results apply to other populations.

Finally, while we observe ancestry-associated differences in IL-10 signaling and fibroblast activation, our study is not designed to disentangle genetic from environmental or sociocultural contributors to these patterns.^44–46^ Self-reported ancestry reflects a complex combination of genetic background and lived experiences that can shape immune and inflammatory states.^27,47,48^ Our findings should therefore be interpreted as correlations that generate hypotheses rather than definitive causal mechanisms.

Overall, our work identifies IL-10 signaling and anti-inflammatory macrophage activation as important modulators of fibroblast activation in response to oxidative stress. This suggests that differences in these pathways across self-reported ancestry groups may contribute to variability in fibrosis-associated phenotypes. The PEG-based hydrogel platform described here recapitulates key features of the fibroblast response to injury. This model can be used to incorporate circulating cells and serum from individual donors to assess fibrosis risk or evaluate candidate therapeutics. Future studies leveraging this system to interrogate additional immune subsets, demographic groups, and pharmacologic modulators, such as inhibition of TGF-β1 signaling to further elucidate the role of TGF-β1. This has the potential to reveal new strategies to prevent or treat fibrosis following injury.

## Methods

### Isolation of serum and monocytes from recruited participants

Participants were recruited in accordance with the Declaration of Helsinki and with IRB approval (2047810-4) from the University of Maryland College Park (UMCP) IRB, College Park, MD, KIRB 0223. Thirty-two individuals aged 18 to 45 of self-reported Black/African or White/European descent were recruited using fliers. Recruited individuals provided informed consent before participation. Participants filled out a survey providing information on age, sex, self-reported European or African ancestry, and other demographic information. A phlebotomist drew blood (45 mL) from each participant. Peripheral blood mononuclear cells (PBMCs) were isolated from whole blood using BD Vacutainer® CPT™ Mononuclear Cell Preparation Tubes (BD Biosciences, according to the manufacturer’s instructions. Monocytes were isolated from the peripheral blood mononuclear cell (PBMC) fraction using pan-monocyte negative selection with QuadroMACS™ separator (Miltenyi). Cells were cryopreserved using CryoStor® CS10 (STEMCELL Technologies). Serum was also obtained and stored at −80°C.

### Serum and monocytes for NanoString and mass spectrometry analysis

Aliquots from thirty-two donors were sent for mass spectrometry analysis (Poochon Scientific). RNA from Monocytes was isolated using an RNeasy Minikit (Qiagen,). RNA from twenty-four donors was sent for NanosString analysis using an nCounter® inflammation panel. Monocytes and serum from the same donors were used for NanoString and mass spectrometry analysis, respectively. Macrophages were taken from twelve donors for hydrogel experiments. Of these twelve donor macrophages, nine were also used for NanoString analysis.

### Cell culture

Primary dermal fibroblasts were purchased from iXCells Biotechnologies. They were obtained from three healthy women of European ancestry aged 20, 25, and 34, and one healthy woman of African ancestry aged 31. Fibroblasts were cultured in fibroblast growth medium supplemented with 10% fetal bovine serum (FBS) and 1× antibiotic–antimycotic (Thermo Fisher) and were used between passages 3 and 5. Human peripheral blood mononuclear cells (PBMCs) were purchased from iXCells Biotechnologies from four healthy Caucasian females aged 23, 24, 27, and 34 years. PBMCs were cultured in RPMI 1640 medium supplemented with 10% FBS and 1× antibiotic–antimycotic (Thermo Fisher Scientific). Monocytes isolated from recruited donors were cultured in donor medium consisting of 5% autologous donor serum and 5% FBS in fibroblast growth medium supplemented with 1× antibiotic–antimycotic (iXCells Biotechnologies). All cells were maintained at 37 °C in a humidified incubator with 5% CO_2_.

### Macrophage polarization

PBMCs purchased from four individuals (iXCells Biotechnologies) or thawed monocytes from twelve recruited participants were plated at 3×10^6^ cells/well on 6-well plates. After 24 hours, cells were rinsed twice in phosphate-buffered saline without calcium and magnesium (1X PBS, Fisher Scientific), and PBMC media was added. After another 48 hours, media was replaced with PBMC media containing macrophage colony-stimulating factor (20 ng/mL M-CSF, Thermo Fisher). Cells were cultured in this medium for 72 hours. Following M-CSF treatment, adherent macrophages were cultured for 48 hours in PBMC media, PBMC media with interleukin-4 (20 ng/mL, Prospec), or PBMC media with LPS (100 ng/mL, Prospec) and IFNγ (10 ng/mL, Prospec).

### Peptide conjugation

Polyethylene glycol was conjugated to the arginine-glycine-aspartate-seri ne peptide (RGDS, MW = 433 g/mol, ThermoFisher Scientific) and GGGPQGIWGQGK (PQ peptide, MW = 1141 g/mol; Genscript) by amine substitution reactions. RGDS and acrylate-PEG-succinimidyl valerate (MW = 3400 g/mol; Laysan Bio) were mixed at a 1.2:1 molar ratio in HEPBS (20 mM at pH 8.5) and titrated to pH 8.0 using NaOH (0.1 M). The mixture was wrapped in foil to protect from light and left overnight at 4°C with constant agitation. The resulting acrylate-PEG-RGDS was dialyzed using a 3.5 kDa MWCO cellulose membrane (Spectrum Laboratories) and stored at −80°C overnight. The frozen mixture was then lyophilized and stored at −80°C until use. PQ conjugation was performed by mixing PQ and acrylate-PEG-succinimidyl valerate at a 1:2 molar ratio in HEPBS, with titration to pH 8.0 using 0.1 M NaOH. The mixture was left overnight at 4°C with constant agitation and protection from light, followed by dialysis with a 6-8 kDa MWCO cellulose membrane (Spectrum Laboratories). The product was then frozen overnight at −80°C, lyophilized, and stored at −80°C until use.

### Cell encapsulation

Hydrogels (5 µL) were formed using PEG-RGDS (3.5 mM) and PEG-PQ-PEG (6% w/v) dissolved in HEPES buffered saline (10 mM HEPES, 100 mM NaCl at pH 7.4, and 1.5% TEOA), the photoinitiator eosin Y (10 µM), and N-Vinyl-2-pyrrolidone (0.35%, Fisher Scientific). Each 5µL hydrogel mixture contained 5 × 10^4^ or 7.5 × 10^4^ dermal fibroblasts pooled from one donor of African ancestry and one donor of European ancestry, with or without the addition of macrophages at a 1:10 ratio from a single donor (5 × 10^3^ or 7.5 × 10^3^ macrophages). Macrophage donors were not matched to the fibroblast donors. Encapsulated cells (5 µL) were placed on a polydimethylsiloxane slab between two polydimethylsiloxane spacers approximately 300µm in height. A methacrylated coverslip was placed on top of the encapsulated cells and the two spacers, followed by exposure to white light for 60 seconds. The coverslip was then inverted and placed in a 24-well plate. Fibroblast media was gently pipetted into the well, with 10% fetal bovine serum and 1X antibiotic-antimycotic (iXCells Biotechnologies).

### Hydrogen peroxide and cytokine treatment

Cells encapsulated in hydrogels were cultured in fibroblast media or donor media for 24 hours with or without TGF-β1 (20 ng/mL, R&D Systems), followed by washing twice in 1X PBS. Experiments with macrophages from vendors used fibroblast media, while experiments with cells from recruited donors used donor media. The cells were then cultured in media 24 hours with or without 20 ng/mL TGF-β1 and with or without H_2_O_2_ (125 µM or 250 µM). This was followed by two washes in 1X PBS. Cells for ‘0 day’ time-points were then isolated for additional assays. Cells for ‘3 day’ and ‘7 day’ time-points were instead cultured for an additional 3 days or 7 days, respectively, in media with or without TGF-β1 (20 ng/mL), and without H_2_O_2_. Similar treatments were performed, replacing TGF-β1 with Stattic (20µM, R&D Systems).

### Immunostaining and live/dead stain

Encapsulated cells were rinsed twice with 1X PBS at room temperature. The cells were then fixed in 4% paraformaldehyde for 45 minutes at room temperature in 24-well tissue culture plates (Fisher Scientific), followed by three washes of 5 minutes each with 1X tris-buffered saline (TBS). The cell membrane was permeabilized using 0.25% Triton-X for 45 minutes at room temperature, followed by three 10-minute TBS washes. Encapsulated cells were blocked with 5% donkey serum overnight at 4°C and then washed three times with TBS for 5 minutes per wash. Samples were incubated overnight at 4°C with primary antibodies in 0.5% donkey serum. Primary antibodies were used at the following concentrations: 1:350 CD90 (cat. PA5-19153, ThermoFisher Scientific), 1:500 alpha-smooth muscle actin (cat. 14-9760-82, ThermoFisher Scientific), 1:1000 collagen 3 (cat. ab184993, abcam), 1:750 Ki67 (cat. 14-5699-80, ThermoFisher Scientific), 1:350 p16INK4a / CDKN2A (cat. A43F5779-SP, R&D Systems), and 1:1000 active caspase-3 (AF835-SP, R&D Systems).

Samples were washed at 4°C five times with Tween in TBS (0.01%) for 90 to 120 minutes for each wash, followed by a final wash with TBS for 90 to 120 minutes. Samples were then incubated with secondary antibodies overnight at 4°C. The following secondary antibodies were used at a 1:200 concentration: Alexa Fluor 488 donkey anti-rabbit (cat. A-21206, ThermoFisher Scientific), Alexa Fluor 555 donkey anti-mouse (cat. A-31570, ThermoFisher Scientific), and Alexa Fluor 647 donkey anti-goat (cat. A-32849, ThermoFisher Scientific). The encapsulated cells were then washed in TBS for 3 hours at 4°C, stained with DAPI (2 μM, cat. 10236276001; Sigma Aldrich) for at least 3 hours at 4°C, and washed twice with TBS at 4°C for 5 minutes per wash.

### Imaging and image analysis

Hydrogels were imaged using the BC43 benchtop confocal microscope (Andor) using the blue, green, red, and far red channels. Cells were identified using DAPI. Identical gain and exposure were used across samples within the same experiment. Images within the hydrogel were taken as Z-stacks with a step unit of 1.5 µm and a range of 40 µm. Image planes were selected in a random and unbiased manner. Individual planes were imported into Imaris, and a Z-projection was made by combining slices. Cells were identified using the Imaris spot analysis. The Imaris machine learning algorithm was then trained to identify positive cells. The output of the algorithm was checked for accuracy and then applied to the Z-projections. The number of positive and negative cells was then quantified.

### Reverse transcriptase quantitative polymerase chain reaction

Encapsulated cells were rinsed twice with 1X PBS at room temperature. The gels were then transferred to microcentrifuge tubes and frozen in liquid nitrogen. RLT lysis buffer from the RNeasy Mini Kit (Qiagen) was added to the gels, followed by homogenization using a pestle. RNA quantification was performed using a NanoDrop spectrophotometer. RNA was reverse transcribed using iScript™ Reverse Transcription Supermix Kit (Bio-Rad). cDNA was amplified with iTaq™ Universal SYBR® Green Supermix (Bio-Rad). Forward and reverse primers for *COL3A1, IL10, PPIA*, and *TGFB1* were used in accordance with manufacturer instructions (IDT, see the table below). Cycling parameters were 40 cycles of: 95°C for 25 seconds for DNA denaturation and polymerase activation, 95°C for 2 seconds for denaturation, and 60°C for 25 seconds for annealing and extension. Relative gene expression was calculated using the 2^(-ΔΔCT) method, with *PPIA* as the internal control.

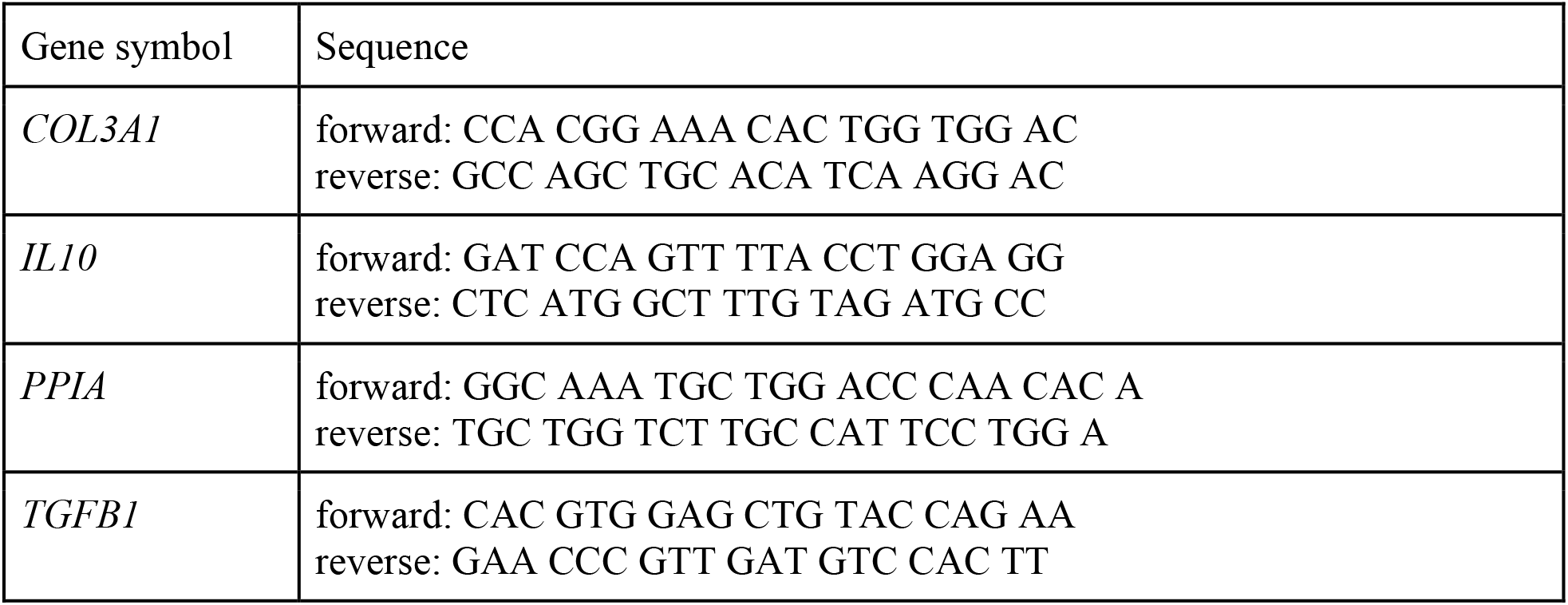

## Statistical analysis

GraphPad Prism was used for statistical quantification and assessment. Following tests for normality, statistical significance was determined using a Kruskal-Wallis test with Dunn’s post-hoc test, with p < 0.05 considered statistically significant. Four biological replicates for experiments with PBMCs and dermal fibroblasts purchased from vendors. Six biological replicates per European or African ancestry groups were used for experiments with monocytes isolated from recruited individuals. Sixteen biological replicates per ancestry group were used for NanoString and mass spectroscopy analysis.

Raw RCC files generated by the NanoString platform were processed using both the nSolver software suite and custom R workflows. Quality control was performed with the NanoString package by evaluating positive and negative control probe performance. Detection thresholds were defined as two times the median, and samples with background signal exceeding this threshold were flagged as failing quality control. Data were normalized using the means of each housekeeping gene before downstream analysis. Differential gene expression analysis was conducted using the limma package, fitting gene-wise linear models to normalized expression data with empirical Bayes variance moderation. P-values were adjusted for multiple testing using the Benjamini-Hochberg false discovery rate (FDR) procedure. Statistical significance required an adjusted p-value of less than 0.05 as well as a minimum absolute log2 fold-change of 0.5

Proteomic data were provided by Poochon Scientific as Excel files containing normalized and quantified protein intensities generated by mass spectrometry. All initial preprocessing, normalization, and quality control were performed by Poochon Scientific. Downstream statistical analyses were conducted in R, where differential protein abundance was assessed using the limma package by fitting gene-wise linear models with empirical Bayes variance moderation. P-values were adjusted for multiple testing using the Benjamini-Hochberg false discovery rate (FDR) procedure. Statistical significance was defined as an adjusted p-value of less than 0.05 together with a minimum absolute log2 fold-change of 1.0. Exploratory gene ontology and pathway enrichment analysis were performed using clusterProfiler with Benjamini-Hochberg FDR correction, using the default background of GO-annotated human genes.

Data visualization was performed using ggplot2, with selected boxplots generated in GraphPad Prism. All analyses were conducted in R version 4.5.0. Analysis code is available at https://github.com/pabloprd/R35_analysis_and_data_generation/tree/main.

## Data availability

The mass spectrometry proteomics data generated in this study have been deposited in the MassIVE repository under accession number MSV000100385 and are also available via ProteomeXchange with identifier PXD072700. NanoString gene expression data generated in this study have been deposited in the Gene Expression Omnibus (GEO) under accession number GSE313951.

## Code availability

The full code used to perform the data analyses presented in this study is publicly available at the following github repository: https://github.com/pabloprd/R35_analysis_and_data_generation/tree/main. This repository contains all scripts and documentation required to reproduce the RNA-seq and proteomics analyses supporting the results in this paper.

## Acknowledgements

This study was funded by NIGMS R35 MIRA GM147048-05. The funder played no role in study design, data collection, analysis and interpretation of data, or the writing of this manuscript.

## Author contributions

N.O.B. - conceptualization, experimental design, formal analysis, methodology, figure preparation, writing–original draft preparation, writing review and editing. A.M.V. - macrophage differentiation and encapsulation. M.T. - donor recruitment, monocyte and serum collection, and writing methodology. P.P. - NanoString and mass spectroscopy analysis, and writing methodology. K.B.K. - encapsulation and PCR analysis. S.R. - encapsulation and PCR analysis. E.M. - methodology, experimental design, supervision, visualization, writing– original draft supervision and modification, writing-review and editing. All authors reviewed the manuscript.

## Competing interests

The authors declare no competing financial or non-financial interests.

## Notes

https://github.com/pabloprd/R35_analysis_and_data_generation/tree/main

### Competing Interest Statement

The authors have declared no competing interest.

