## Supplemental file for "Ancestry-Linked IL-10 Signaling and Macrophage Activation Modulate Fibroblast Responses to Oxidative Stress in a PEG-Based Microphysiological System"

Supplementary Figures

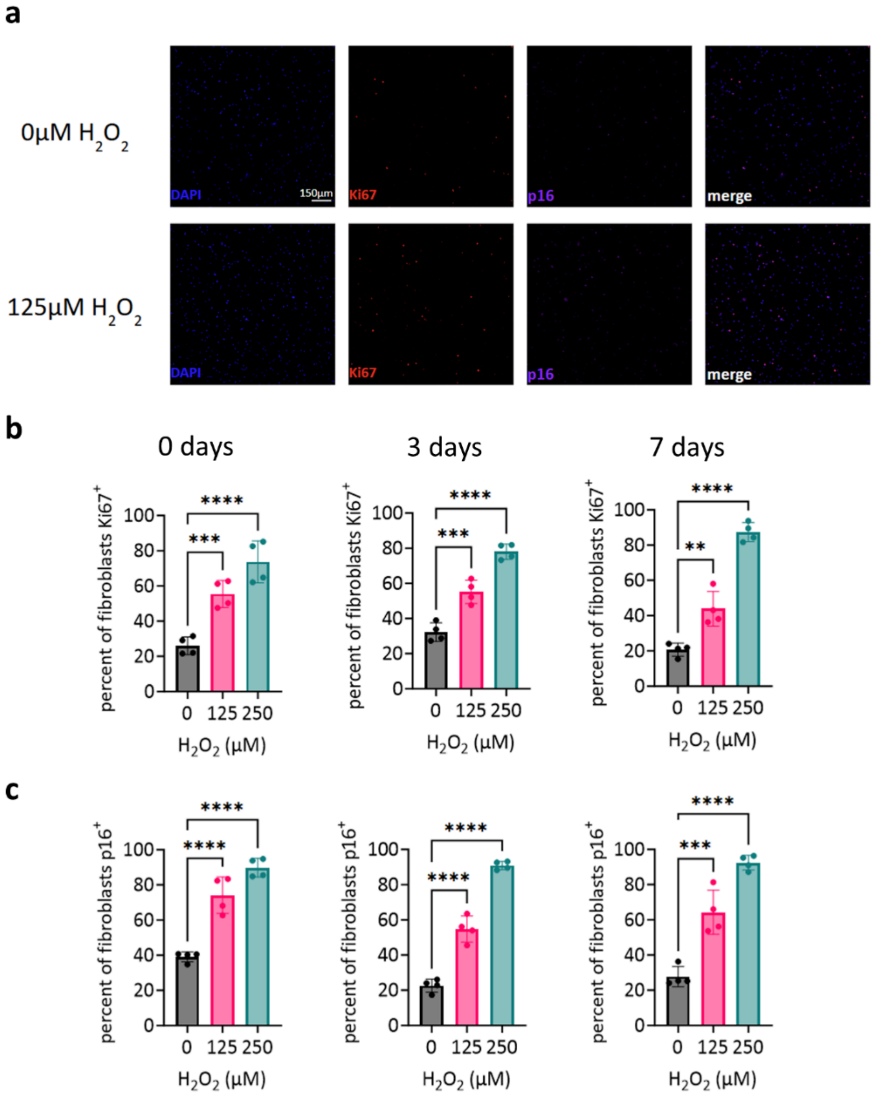


Supplement Figure 1. Oxidative stress dose–response profiles for Ki67 and p16 expression. (a) Representative immunocytochemistry images of fibroblast expression of Ki67 [red], p16 [purple], and nuclei [DAPI in blue] 3 days after $H_2O_2$ treatment. (b) Percent of fibroblasts expressing Ki67, and (c) p16 at 0, 3, and 7 days after $H_2O_2$ treatment. n = 4 biological replicates. Data shown as mean ± standard deviation. The Kruskal-Wallis test was used to calculate p-values and statistical significance. ***p*-value < 0.01; ****p*-value < 0.001, *****p*-value < 0.0001.

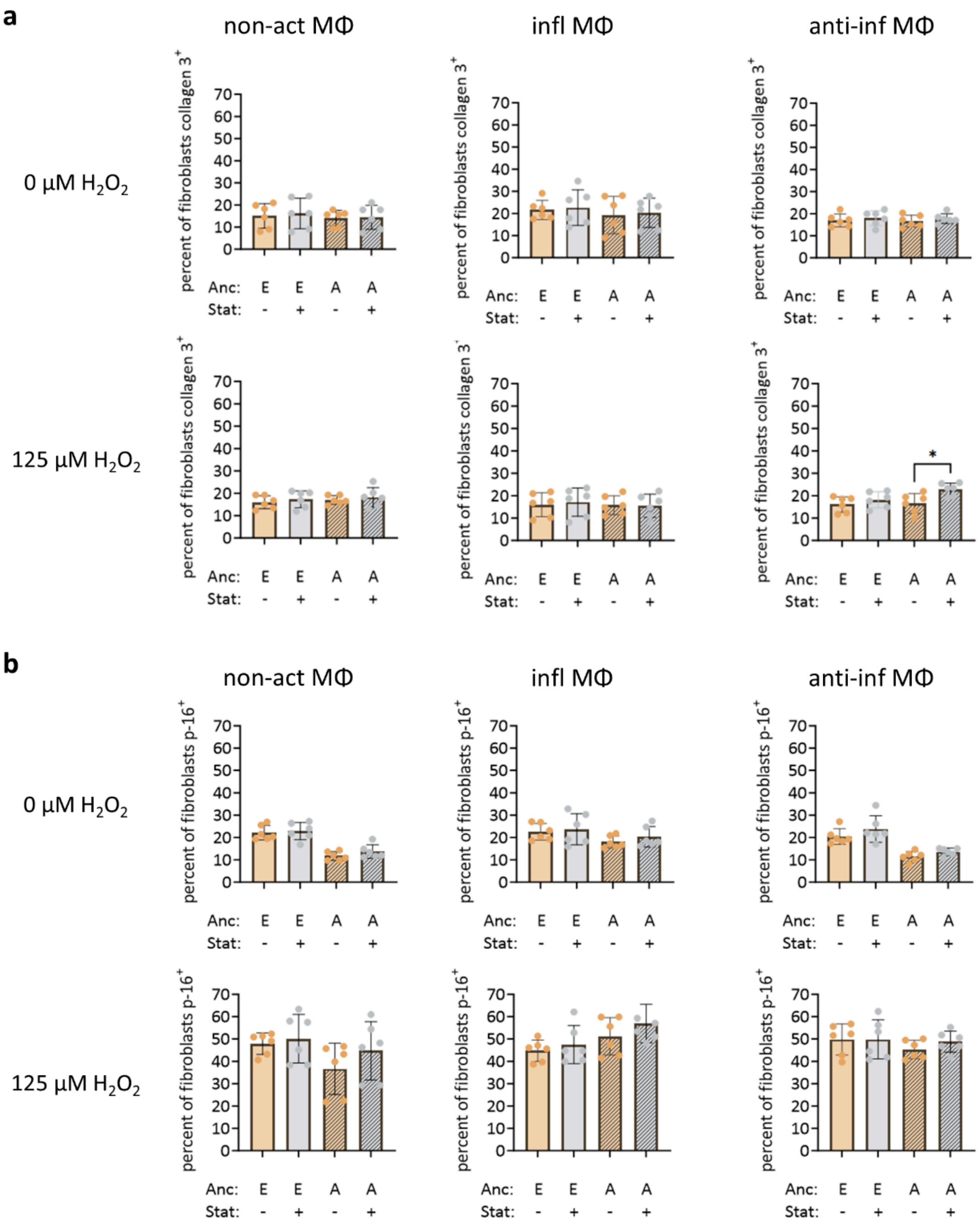


Supplemental Figure 2. IL-10 signaling did not mediate differences in fibroblast expression of collagen 3 and p16. Percentage of fibroblasts expressing (a) collagen 3 and (b) p16 at 7 days after $H_2O_2$ treatment, culture in donor serum, with or without Stattic treatment, and co-encapsulation with unactivated macrophages (non-act Mθ), inflammatory macrophages (inf Mθ), and anti-inflammatory macrophages (anti-inf Mθ). Macrophages and serum are from donors of self-reported European (E) or African (A) ancestry. n = 6 biological replicates per ancestry group. The Kruskal-Wallis test was used to calculate p-values and statistical significance.
